# Structural ensembles based on NMR parameters suggest a complex pathway of ligand binding in human gastrotropin

**DOI:** 10.1101/409383

**Authors:** Zita Harmat, András L. Szabó, Orsolya Toke, Zoltán Gáspári

## Abstract

Gastrotropin, the intracellular carrier of bile salts in the small intestine, binds two ligand molecules simultaneously in its internal cavity. The molecular rearrangements required for ligand entry are not yet fully clear. To improve our understanding of the binding process we combined molecular dynamics simulations with available structural and dynamic NMR parameters. The resulting ensembles reveal two distinct modes of barrel opening with one corresponding to the transition between the *apo* and *holo* states, whereas the other affecting different protein regions in both ligation states. Comparison of the calculated structures with NMR-derived parameters reporting on slow conformational exchange processes suggests that the protein undergoes partial unfolding along a path related to the second mode of the identified barrel opening motion.

## Introduction

Gastrotropin (also known as ileal bile acid-binding protein (I-BABP) or fatty acid-binding protein 6 (FABP6)) [1] is involved in the enterohepatic circulation of bile salts from the liver to the small intestine and back to the liver. This recirculation process ensures that only a small amount of bile salts needs to be synthesised *de novo*. The protein is located in the epithelial cells of the distal small intestine [2-3] and has an important role in cholesterol homeostasis [1,4].

Gastrotropin belongs to the family of intracellular lipid-binding proteins (iLBPs), a group of small, approximately 15-kDa proteins that bind fatty acids, retinoids, cholesterol, and bile salts [5]. Additionally, iLBPs have been shown to have a role in the stimulation of the transcriptional activity of nuclear hormone receptors [6-8]. Among the four main groups of the iLBP family, the subfamily of gastrotropin is unique in the sense that it has the capability of binding two [9-10] or possibly even three [11-12] ligands simultaneously. NMR solution structure of the *apo* form of human gastrotropin (PDB ID: 1O1U) was determined along with the cholyltaurine bound form (PDB ID: 1O1V) [13]. More recently, the structure of the heterotypic doubly-ligated complex of human gastrotropin with glycocholate and glycochenodeoxycholate has been determined [14]. Similarly to other members of the iLBP family, the structure of human I-BABP is composed of a β-barrel formed by ten antiparallel β- strands (A-J) and two α-helices (I-II). The binding cavity of ∼1000 Å^3^ is located inside of the β-barrel [15]. Ligand binding in human gastrotropin exhibits positive cooperativity [9], which has been shown to be governed by the hydroxylation pattern of the bound bile salts [16]. Accordingly, hydrogen bonding networks have been shown to have a key mediatory role in positive binding cooperativity [17]. Besides the observed communication between the two binding sites, di- and trihydroxy bile salts display a site preference upon binding in each other’s presence [18]. As it is apparent from the comparison of *apo* and *holo* human gastrotropin structures (Fig 1), bile salt binding is accompanied by large conformational changes in the E-F and G-H protein regions as well as in the C/D-turn and the proximate helical cap [2,11]. Importantly, NMR relaxation measurements suggest that in the *apo* form, the ground state is in slow exchange with a low-populated ‘invisible’ conformer resembling some structural features of the the ligand-bound form [19]. Intriguingly, residues undergoing a conformational fluctuation on the μs-ms time scale can be grouped into a ’slower’ and a ’faster’ cluster, which appear to be spatially separated. Specifically, while the ‘slower’ cluster involves part of the helical region, the C/D-turn, and the proximate B and D β-strands in the N-terminal half, the ‘faster’ cluster comprises segments of the EFGH protein region in the C-terminal half [19].

**Fig 1:**
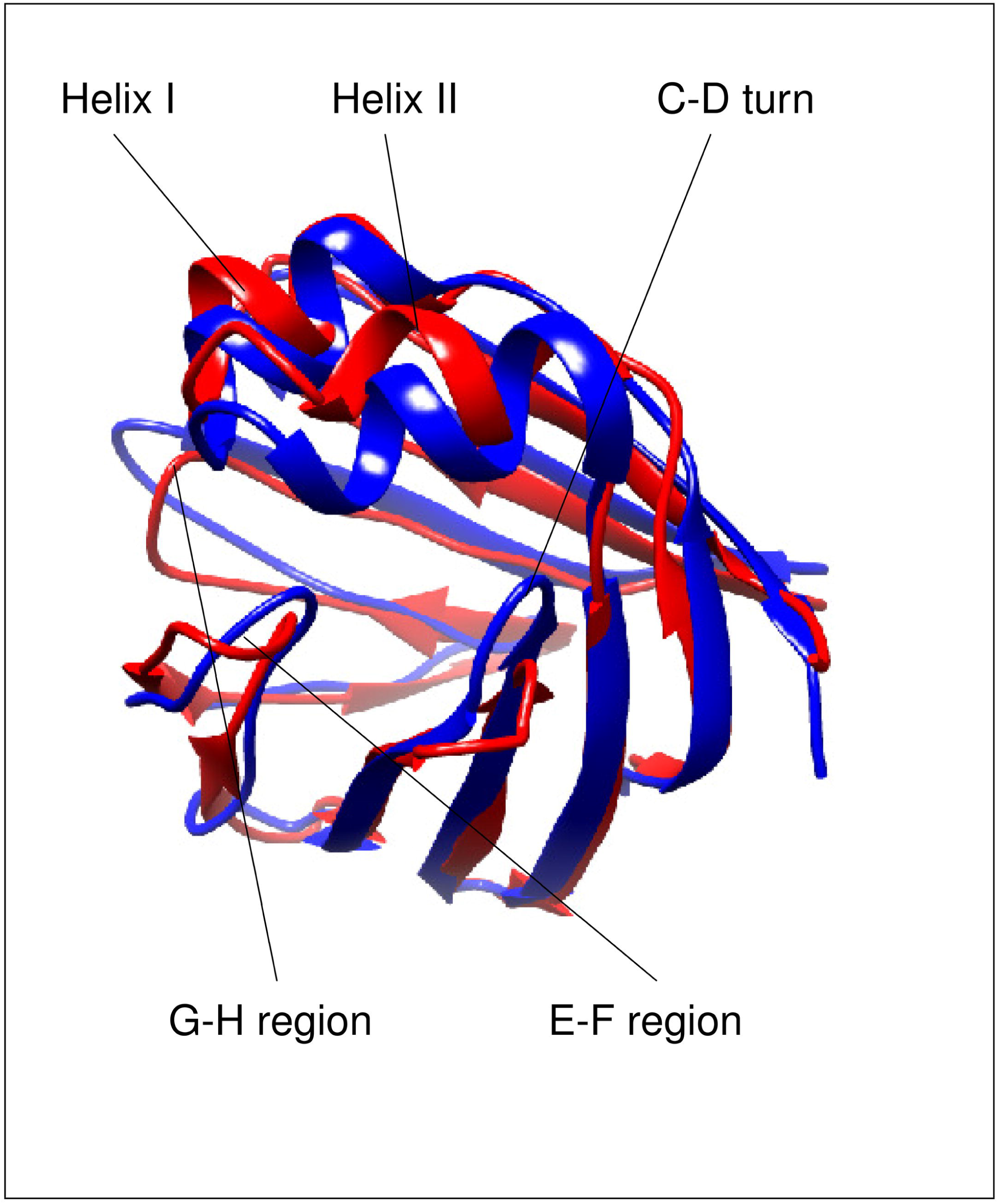
Ribbon representation highlighting the differences between the *apo* (PDB ID: 1O1U model 7) and the *holo* (PDB ID: 2MM3 model 1) form of human gastrotropin. The figure was prepared with UCSF Chimera [45].

As the binding site of gastrotropin is located in the interior of the protein, the mechanism of ligand entry is an important issue to be investigated. The most widely accepted scenario for the protein family is formulated in the ‘portal hypothesis’, stating that access of ligands to the protein interior is governed by the C/D and E/F-turn regions together with the C-terminal part of helix II [20-22]. Based on NMR structural and dynamic studies, a conformational selection mechanism of ligand binding involving an equilibrium between a closed and a more open protein state has been suggested for both the human ileal [14] and the chicken liver BABP analogues [23]. In line with the NMR spectroscopic analysis of internal motions, molecular dynamics simulations show evidence of correlated motions in human gastrotropin and in the absence of ligands indicate a partial unfolding of the E-F protein region [24].

To improve our understanding of the mediatory role of internal motions in human gastrotropin-bile salt interaction, we generated conformational ensembles consistent with experimentally obtained NMR structural and dynamic data [25] and performed ligand docking to obtain an atomic-level insight into the binding mechanism. Our results reveal different conformational rearrangements in the protein that are suggested to correspond to motions characteristic of different time scales indicating a complex mechanism of bile salt entry.

## Materials and Methods

### Ensemble molecular dynamics simulations with NMR restraints

Molecular dynamics calculations were performed using GROMACS version 4.5.5. [26-27] modified to handle S^2^ order parameters as well as pairwise averaging of NOE distance restraints over replicas [28], as proposed for the MUMO (Minimal Under-restraining Minimal Over-restraining) approach [29]. The OPLS-AA force field [30] and the TIP3P water model [31] was used for all molecular dynamics simulations described below.

For modeling the *apo* structure of gastrotropin, we chose model 7 of PDB entry 1O1U [13] based on its highest PRIDE-NMR score [32] among the deposited models. As an initial model of the *holo* structure we used model 1 of the PDB entry 2MM3. Ligand topologies for glycocholic acid (GCA, PDB ligand ID: GCH) and glycochenodeoxycholic acid (GCDA, PDB ligand ID: CHO) were generated with the TopolGen script and corrected manually for atom types where necessary as well as with an in-house Perl script to reassign hydrogen atoms to the charge groups defined by the heavy atoms they are connected to.

NOE restraints were only available for the *holo* protein (PDB ID: 2MM3). For the *apo* form, we used restraints from the 2MM3 list that were unviolated in the deposited 1O1U structure as checked with the CoNSEnsX server. Restraints were modified by the removal of stereospecificity and rounding the restrained distance up to the next integer Å, creating 1 Å wide ‘bins’ from 4 to 10 Å.

Chemical shifts for the *apo* structure were obtained from BMRB (BMRB ID: 19843) and for the *holo* structure directly from the authors. S^2^ values for the *apo* and *holo* structures measured at 283, 291, 298, and 313 K were taken from [19].

After generating a topology using the OPLS-AA force field and TIP3P water model, the molecule was put into a cubic box, followed by energy minimization with conjugate gradient method for 5000 number of steps with 0.001 ps step length. The maximum force was set to 200. In the next step, the molecule was solvated and then one of the water molecules was replaced by a Na^+^ ion to ensure the neutrality of the system. After that, another energy minimization was performed using the same parameters, but including the water molecules. In the last step, a short MD simulation was performed using position restraints of 1000 kJ mol^-1^ nm^-2^ on the heavy atoms of the protein for 2500 steps with 0.002 ps step size using the LINCS algorithm [33].

For the production runs, eight replicas were simulated in parallel with the OpenMPI environment [34]. Backbone S^2^ order parameter restraints were applied on the full ensemble and NOE distance restraints were averaged between neighboring replicas, similar to the MUMO (Minimal Under-restraining, Minimal Over-restraining) protocol [29]. The simulations were performed at four temperatures: 283 K, 291 K, 298 K, and 313 K using S^2^ restraints measured at the corresponding temperatures. With LINCS constraining on bond lengths, a timestep of 2 fs was used to generate runs of 2 ns and 6 ns, totaling 16 and 48 ns for the 8 replicas combined, respectively. Control simulations with the same parametrization but without restraints were also performed. Topology files for the restrained simulations are included in the supplementary material as (File S1) - (File S8).

In order to generate a larger pool of possible conformations in order to further explore the conformational space, molecular dynamics simulations with only one type of restraint, NOE or S^2^, or without any restraints were also performed. Accelerated Molecular Dynamics and short (500 ps) Targeted Molecular Dynamics simulations were also performed on the *apo* structure using the chemical shifts of the *holo* structure and vica versa in order to achieve transition from one form to the other.

### Docking Simulations

Docking calculations were performed on selected structures with the ligands GCA and GCDA using Schrödinger Glide [35]. The binding sites were defined using the ternary complex structure 2MM3. After importing the structure, it was split to separate molecules. As a next step, either GCA or GCDA was merged with the protein and a mesh grid around the ligand was generated with the ’Receptor Grid Generation Tool’ using default settings. Docking of the respective ligands was performed using the ’Ligand docking’ tool with default settings except requiring the inclusion of per-residue interaction scores in the output. To dock the second ligand into the binary complex obtained, the docking result most similar to the pose in the initial 2MM3 structure was merged with the second ligand and used to define the second binding site with the grid generation tool. For each of the four different setups, i.e. GCA, GCDA, GCA+GCDA and GCDA+GCA docking runs, 32 different poses were generated and evaluated. Total energy of the docked complexes was estimated using the MacroModel routine with the OPLS3 force field and water as solvent.

### Data analysis

Correspondence to the experimental parameters was analyzed using the CoNSEnsX webserver [32,36] (for S^2^ order parameters and chemical shift data) as well as in-house Perl scripts (for NOE distance restraints). NOE restraints were evaluated on a per-ensemble basis using r^-6^ averaging both for intramolecular ambiguity and between members of the ensembles.

For the S^2^ order parameter correspondences, MUMO simulations and the original PDB ensembles, the corrected S^2^ values are also displayed. In the correction, those points were excluded from the analysis, which had greater than 0.2 as an absolute value of the difference between the experimental and back-calculated values. For the MUMO simulations, maximum 5 such values were found. All the experimental and back-calculated values are depicted in S1 Fig.

Principal Component Analysis was performed using ProDy [37] and visualized with the NMWIZ module of the program VMD [38]. Structure-based chemical shift calculations were performed with the program SHIFTX2 [39].

The presence or absence of hydrogen bonds in the ensembles was investigated with an in- house Perl program using distance-angle based hydrogen bond identification parameters [40-41].

### Comparing calculated ^15^N chemical shift differences with experimentally derived Δ□ (^15^N) values

For each structure, backbone ^15^N chemical shifts were estimated with Shiftx [39]. For each conformation in the large conformer pool (see above), the absolute value of the difference of the predicted chemical shifts relative to those in each calculated unliganded structure in the MUMO ensembles was calculated. These differences were then compared to experimental Δ □ (^15^N) data derived from CPMG relaxation dispersion NMR measurements for each residue for which it was available [19]. Both correlation and RMSD measures were calculated after normalization to the 0-1 range. As there are Δ □ values available for three temperatures and the conformational pool is of a heterogeneous source with no well-defined temperature, the correlation and RMSD values were calculated for all three temperatures and then were averaged for each structure investigated. The structures with highest correlation and lowest RMSD values were selected for analysis.

## Results and Discussion

### The generated ensembles reflect experimental parameters

According to the expectations, experimental S^2^ parameters are generally better reflected in the MUMO generated ensembles than in the PDB ensembles or the unrestrained ensembles (Table 1). Interestingly, the MUMO ensemble of the *apo* protein calculated with the S^2^ parameters of 283 K and the MUMO ensemble of the ternary complex calculated with the S^2^ parameters of 291 K corresponds only moderately to these data, while all other restrained ensembles show good correspondence. The reason for this is the presence of some extremely low (< 0.3) experimental S^2^ values, located mostly in turn regions, not reflected in the simulations. Plots for the experimental and the back-calculated S^2^ values are depicted in S1 Fig.

**Table 1:** RMSD values, S2 parameters, NOE correspondence, amide N and H chemical shift correlations.

|  | Ensemble size | Temperature (K) | Backbone RMSD (Å) | Backbone S<sup>2</sup> correlation | Corrected backbone S<sup>2</sup> correlation | Percentage of violated NOE restraints (r<sup>-6</sup> > 0.5Å) | Chemical shift correlation |  |
| --- | --- | --- | --- | --- | --- | --- | --- | --- |
|  |  |  |  |  |  |  | amide N | amide H |
| 1O1U pdb | 10 | 305 | 0.66+-0 | 0.502<sup>a</sup> | 0.857 | 0.00 % | 0.709 | 0.596 |
| *apo* MUMO | 168 | 283 | 1.01+-0.03 | 0.675 | 0.873 | 0.29 % | **0.832** | **0.72** |
| *apo* MUMO | 168 | 291 | 1.12+-0.04 | 0.8 | 0.841 | 0.29 % | **0.828** | **0.729** |
| *apo* MUMO | 168 | 298 | 1.27+-0.06 | 0.797 | 0.883 | 0.12 % | **0.834** | **0.738** |
| *apo* MUMO | 168 | 313 | 1.45+-0.04 | 0.963 | 0.972 | - | **0.834** | **0.705** |
| *apo* unrestrained | 48 | 283 | 1.46+-0.09 | 0.307 | - | - | **0.827** | **0.724** |
| *apo* unrestrained | 48 | 291 | 1.45+-0.05 | 0.351 | - | - | **0.832** | **0.703** |
| *apo* unrestrained | 48 | 298 | 1.53+-0.03 | 0.301 | - | - | **0.833** | **0.731** |
| *apo* unrestrained | 48 | 313 | 1.5+-0.07 | 0.6 | - | - | **0.829** | **0.733** |
| 2MM3 pdb | 10 | 293 | 0.52+-0 | 0.297<sup>b</sup> | 0.315 | 0.00 % | 0.773 | 0.593 |
| *holo* MUMO | 168 | 283 | 1.28+-0.03 | 0.726 | 0.756 | 0.41 % | **0.784** | *0.548* |
| *holo* MUMO | 168 | 291 | 1.31+-0.07 | 0.544 | 0.731 | 0.46 % | **0.78** | *0.589* |
| *holo* MUMO | 168 | 298 | 1.46+-0.15 | 0.884 | 0.930 | 0.36 % | *0.767* | *0.552* |
| *holo* MUMO | 168 | 313 | 1.5+-0.08 | 0.724 | 0.870 | 0.36 % | *0.758* | *0.551* |
| *holo* unrestrained | 48 | 283 | 1.92+-0.64 | 0.073 | - | - | **0.782** | *0.531* |
| *holo* unrestrained | 48 | 291 | 2.34+-0.65 | 0.296 | - | - | **0.777** | *0.537* |
| *holo* unrestrained | 48 | 298 | 1.82+-0.07 | 0.484 | - | - | **0.772** | *0.493* |
| *holo* unrestrained | 48 | 313 | 2.59+-2.09 | 0.272 | - | - | **0.775** | *0.544* |

a: $S^2$ data of 313 K (highest correlation from the 4 datasets)
b: $S^2$ data of 298 K (highest correlation from the 4 datasets)

The number of ensemble-calculated NOE violations are below 0.5 percent in each of the MUMO ensembles, despite the clearly higher global RMSD values of the MUMO ensembles than those for the original PDB ensembles. Correspondence to the amide N and H chemical shifts are in the same range for the MUMO and the original PDB ensembles.

### Gastrotropin ensembles reveal two distinct modes of barrel opening

Principal component analysis (PCA) of the ensembles suggests the presence of two kinds of modes, both corresponding to the opening of the barrel structure, termed ‘Type I’ and Type II’ openings below. Type I opening clearly separates the *apo* and *holo* structures along PC1, corresponding to the opening of the barrel between strands F and G. Viewing the structure from the direction of the helices, this *apo* to *holo* structural change can be described as a clockwise rotation of the E/F- and G/H-turns accompanied by a lower amplitude counterclockwise rotation of the C/D-turn and helix-II, resulting in the appearance of a large aperture between the E/F- and G/H-turns at the ‘top’ of the barrel. PC2 or Type II opening, in contrast, primarily affects helix-I and the CD-turn, most prominently resulting in the widening of the interhelical gap and the appearance of an opening between strands D and E.

It is notable that the Type II opening motion occurs in both the *apo* and the *holo* structures. At higher temperatures, the ensembles occupy a larger region of the conformational space along this particular opening mode (S4 Fig). It should also be noted that this kind of opening is also compatible with the portal hypothesis as described in detail for human liver fatty acid binding protein [42-43].

The structural changes can be described in more detail by measuring distances between selected amino acids. In S2 Fig A the correlation of Cα distances with each of the two motional modes is plotted for each amino acid pair as a matrix. On the basis of the correlation of these Cα-Cα distances, the most mobile regions corresponding to Type I opening are near the termini and at the D/E-turn region including β strand E itself. Regarding Type II motion, the regions around amino acid 50 (β strand C) and 35 (linker between helix II and β strand B) appear to be the most flexible together moving segments.

Cα-Cα distances displaying the best correlation with Type I opening are between residues 47-69, 48-69, 60-69, 61-69, 62-69, 63-69 (corr. -0.96) as well as 66-70, 67-70 (corr. 0.96). Regarding Type II motion, the best correlated Cα distances are between residues 16-58, 17- 58, and 18-58 (corr. -0.92). The listed Cα atom-atom distances are mapped on the structure in S2 Fig B-C. Our results suggest that the increasing distance between β strands C and E or D and E are correlated with Type I opening, and the increasing distance of helix I from the top of the barrel (around the C/D.turn) correlates with Type II opening. Note that the residues displaying the largest displacement relative to the average structure do not necessarily coincide with the ones exhibiting the largest changes in interatomic distances.

While the experimentally determined *apo* structure (1O1U) is clustered with the dynamic ensembles for the *apo* state, the NMR structure of the ternary complex (2MM3) is located between the *apo* and *holo* ensembles. This indicates that our ensembles of the complex state exhibit a more pronounced opening along PC1 than the structure obtained by conventional NMR calculations. The phenomenon that dynamic ensembles, corresponding reasonably well to experimental data ’magnify’ the differences between different states has been observed in previous works [44] and is likely a consequence of the ensemble-based treatment of NOE restraints allowing more conformational freedom than conventional structure calculations, as well as the different balance between the force field and experimental restraints than in conventional structure calculation methods.

The *apo* and *holo* states exhibit characteristic differences in their hydrogen bond pattern as well (Fig 2B), (Table 2) and (S1 Table). As shown previously, hydrogen bonds form an extensive network in human I-BABP [17,19]. According to our calculations, the most significant differences in hydrogen bond occurence include the formation of one and breaking of two intrastrand hydrogen bonds upon transition from the *apo* to the *holo* state, consistent with a specific mode of barrel opening between strands E and F. Interestingly, hydrogen bonds with ligands (purple lines on Fig 2B) are present only in a few conformations, which may indicate a loose ligand binding as a result of dynamically changing hydrogen bonds. Notable are the hydrogen bonds of Thr73, where the γ1 OH group forms an intraresidue hydrogen bond in the *holo* state that is not present in the *apo* form. This particular residue in the E/F-turn has been suggested to have a key role in a conformational selection mechanism of ligand binding together with proximate residues in the EFGH region of human I-BABP [14].

**Fig 2:**
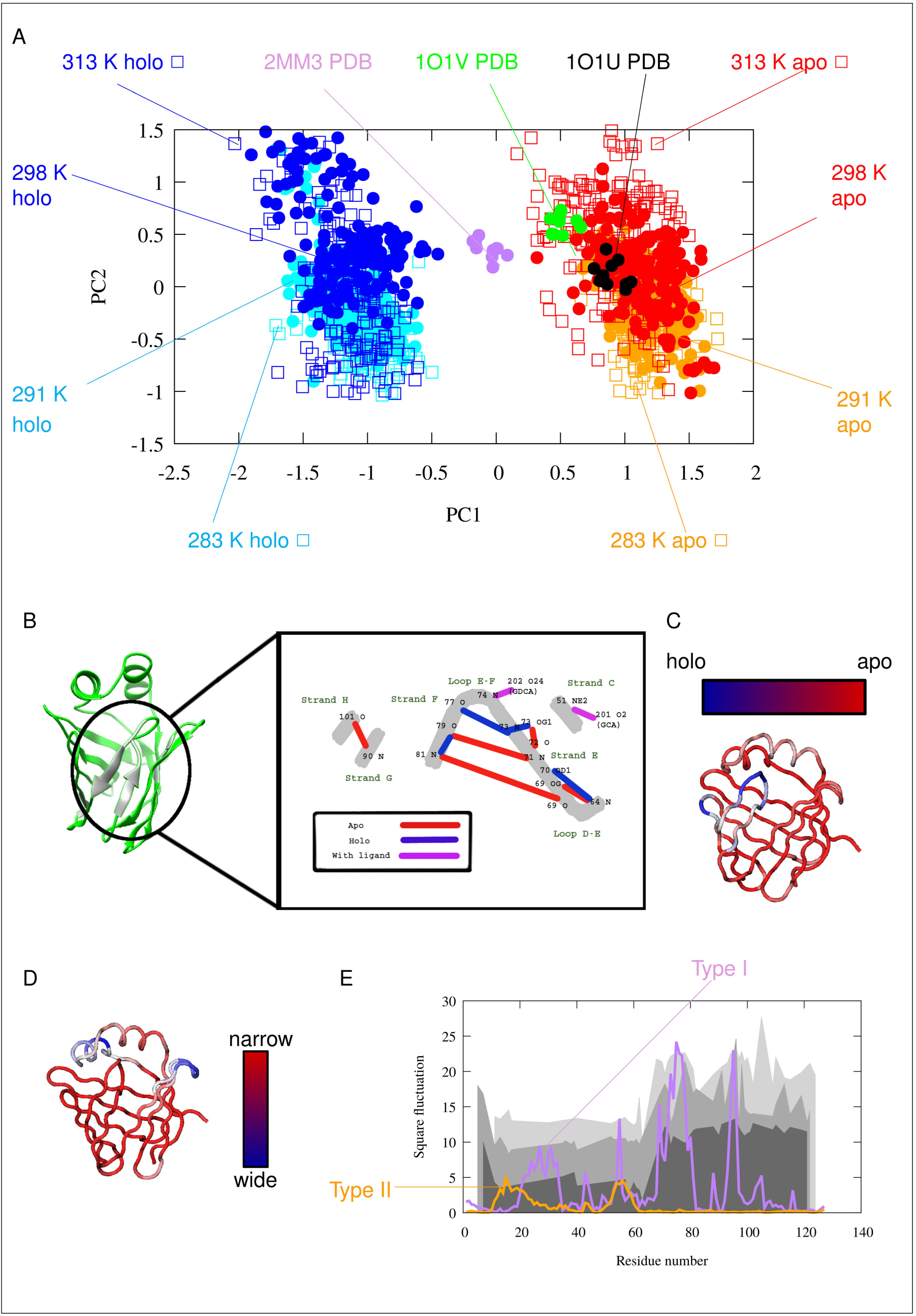
Description of the Type I and Tye II motions. (A) PCA (Principal Component Analysis) scatter plot of the simulated and experimentally determined conformer pool. (B) Hydrogen bonds with the largest changes between the *apo* and *holo* states according to the MUMO ensembles. Black numbers denote amino acid residues, black letters denote atoms, secondary structure elements are labeled with green letters. Red lines represent H- bonds characteristic of the *apo* form, blue lines represent those formed mainly in the *holo* form and purple lines indicate H-bonds between amino acids and the ligands. Note the central role of Thr73 in the hydrogen bond network. (C) Structural movements along PC1: barrel opening (D) Structural movements along PC2 (E) Square fluctuation of Cα atoms in the two PCA modes: PC1 (Type I motion, purple), PC2 (Type II motion, orange). The previously measured experimental k_ex_ values indicating two distinct clusters of residues involved in slow conformational exchange processes are depicted as different gray areas corresponding to the three different temperatures (283 K, 287 K, 291 K) of the measurements. As only about 30-40 amino acids have displayed ms timescale motion with measurable k_ex_ values [19], a continuous depiction is used to guide the eye to highlight the regional differences.

**Table 2:** List of the most significantly changing hydrogen bonds in the course of the MUMO simulations.

| Bond (amino acid number and | Protein state | Donor region | Acceptor |
| --- | --- | --- | --- |

| atom identifier) |  |  | region |
| --- | --- | --- | --- |
| 64 N – 69 OG | *apo* | D-E | E |
| 64 N – 70 OD1 | *holo* | D-E | E |
| 71 N – 79 O | *apo* | E | F |
| 73 N – 73 OG1 | *holo* | E-F | E-F |
| 73 N – 77 O | *holo* | E-F | E-F |
| 73 OG1 – 72 O | *apo* | E-F | E |
| 81 N – 69 O | *apo* | F | E |
| 81 N – 79 O | Mostly *holo* | F | F |
| 90 N – 101 O | mostly *apo* | G | H |

### Residues involved in the two opening modes coincide with different exchange rates along the sequence as determined by NMR

Comparing the regions affected by the motions with NMR-derived conformational exchange data, it is apparent that there is a coincidence of the region affected by Type II opening and the the NMR-reported ’slow’ cluster located in the N-terminal half of the protein [14] (Fig 2E). Although k_ex_ parameters derived from CPMG relaxation dispersion NMR measurements report on a motion occuring on a much slower μs-ms time scale than reflected by the S^2^ restraints used in our simulations, we suggest that the observed Type II barrel opening is related to the slow conformational exchange revealed by NMR relaxation dispersion analysis. Specifically, the fast motions could set the stage for slower, larger-amplitude motions in the protein along a similar opening mode. The structural transition on a different time scale is also consistent with the temperature-dependence of the observed motions, i.e. a more even distribution of conformers along the Type II mode at higher temperatures. Importantly, the presence of fast motion along this mode in both the *apo* and the *holo* states suggests that Type II motions may have a role in both ligand uptake and release.

### The hidden “holo-like” conformation in the apo state is partially unfolded

Previous NMR investigations of human gastrotropin have identified the presence of an invisible state that is in slow exchange with the observable *apo* state [2]. Moreover, it was suggested that this state exhibits *holo*-like structural features [19,14]. In order to get a deeper insight into the nature of this conformer, we generated a pool of conformers and selected structures that might be representative of the higher energy state based on the differences in chemical shifts relative to the *apo* state when compared with the NMR-derived Δ□ (^15^N) values between the ground and higher energy states of IBABP. We note that with the availability of only backbone ^15^N chemical shift differences the structural information on the invisible state remains to be rather limited.

Apparently, the identified conformers with best correspondence to the experimental data are scattered around a large conformational space (Fig 3). Their common characteristics is that they are closer to the *holo* than to the *apo* state, which is in agreement with the previously proposed *holo*-like characteristics of the sparsely populated excited state indicated by NMR dynamic measurements [14]. Importantly, some of the conformers show a more pronounced Type II-like opening than the MUMO ensembles. The principal components in Fig 3. are not direclty corrsponding to those in Fig 2 and the HD3 and HD5 structures, close in the PCA plot, show different degree of Type II-like opening in their helical region. Nevertheless, we consider this aspect the most relevant as Type II opening is clearly identifable in the MUMO ensembles calculated with a substantial amount of experimental data, in contrast to other motions identified in unrestrained simulations only.

**Fig 3:**
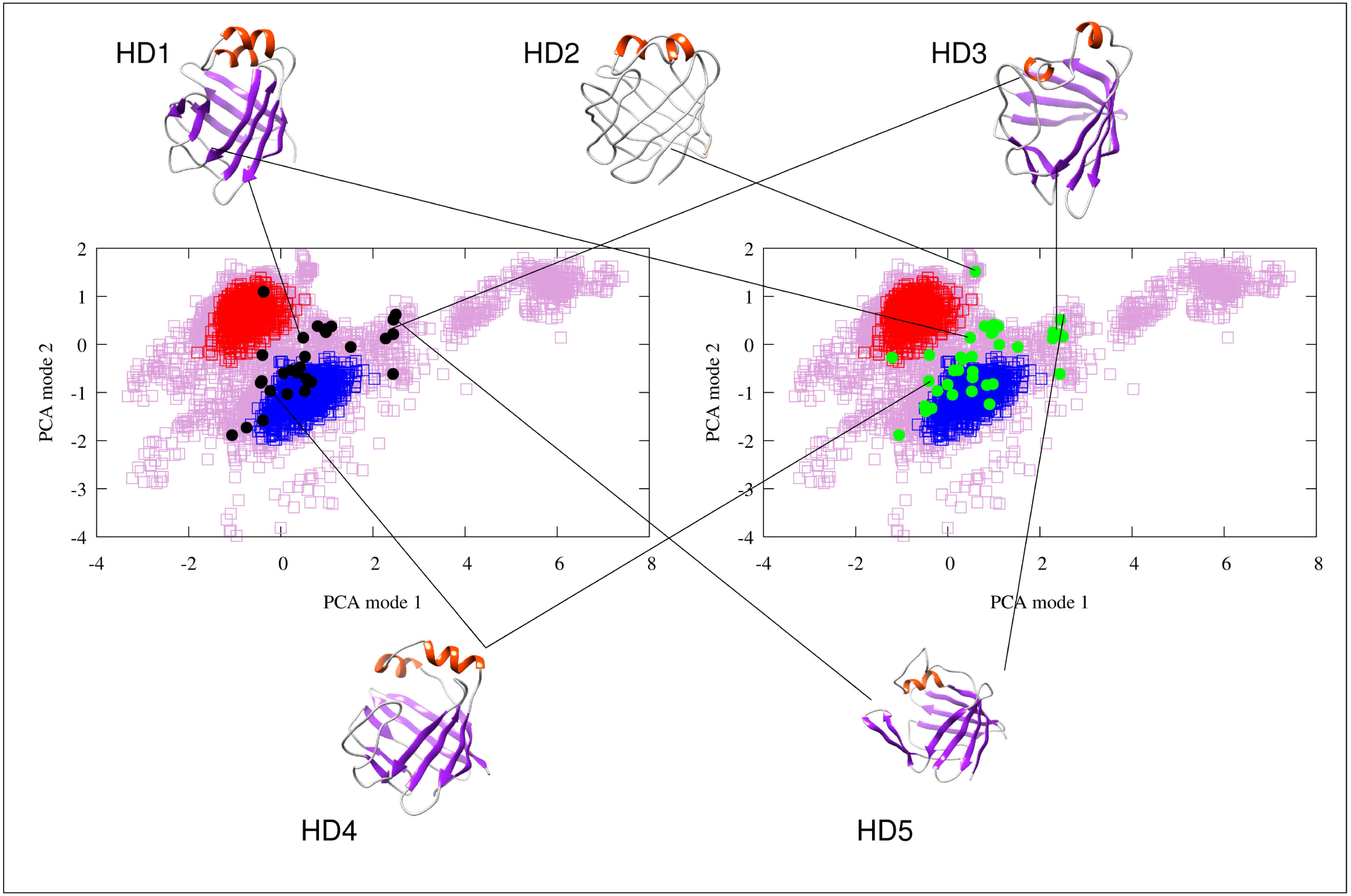
PCA scatter plot of the *apo* MUMO (red dots) and *holo* MUMO (blue dots) ensembles along with the conformer pool (purple hollow squares) used to select the structures best corresponding to the NMR-derived invisible state. Structures with a mean correlation between Δ□ (^15^N) values and calculated chemical shift differences above a threshold of 0.35 are shown with black dots (left panel). Structures with an RMSD between Δ□ (^15^N) values and calculated chemical shift differences lower than 0.00603 are depicted with green dots (right panel). Selected structures are also depicted and linked to their corresponding points in the PCA scatter plots. These hidden conformations are termed HD1- HD5.

As shown in Fig 4, secondary structure of the simulated conformers is diverse around the boundary of the α-helical and β-strand elements. In some structures almost all of the α- helical and β-strand elements are partially unfolded. The structures assumed to be the ’*holo*- like’ *apo* conformations have low helical and β-strand content. The E-F region is the most susceptible to unfolding, in accordance with recent reports by Tőke et al. [24]. Taken together, these observations suggest that the transition from the *apo* to the *holo* state, instead of being a simple physical opening along the shortest route, is rather a complex succession of conformational rearrangements proceeding through a partially unfolded intermediate involving a loosened helical and C/D-turn regions, resembling in part the observed ‘Type II’ mode of motions.

**Fig 4:**
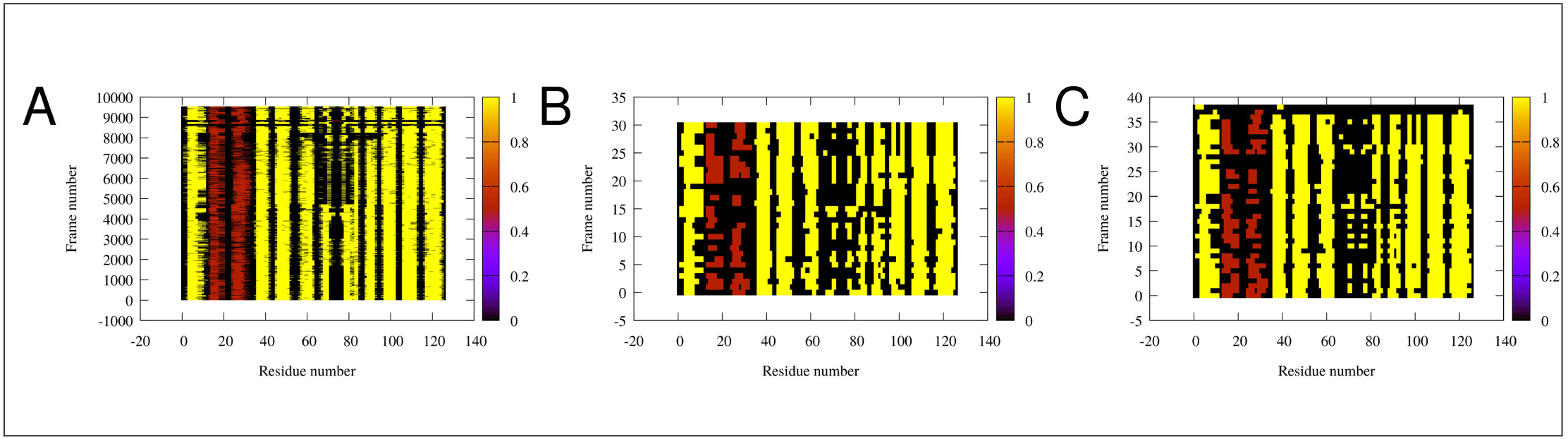
Secondary structure of the conformations inferred from our simulations (rows). Each column represents one amino acid. Extended β-strands are colored yellow, α-helices are brown, the rest of the residues are colored black. (A) All of the conformations. (B) The high correlation conformations (subset of conformations of panel A). (C) Conformations with lowest RMSD (another subset of panel A). The analysis was performed with DSSPCont [46]. Note the shortening of secondary structure elements in some structures, especially in C) and D).

### Docking simulations support cooperativity of ligand binding

In order to further characterize the mechanism of ligand binding, we performed docking simulations into selected structures obtained in our calculations.

Based on the PCA analysis, four structures were selected representing extreme states along Type I and Type II opening, respectively. Additional three structures, regarded as the best models of the invisible state in slow exchange with the *apo* form were also included.

In general, the most favorable complexes were obtained when GCA was docked first, followed by the docking of GCDA (box diagrams in Fig 5). This scenario did not result in a successful ternary complex for only one of the proposed hidden structures, HD1, corresponding to an intermediate position between the apo and holo ensembles along Type I opening. Ligand binding provides the highest stabilization for the partially unfolded structures corresponding to a larger opening along a motion resembling Type II opening. Comparing the relative estimated energies of the corresponding apo structures (colored bars in Fig 5) suggests a complex energetic landscape where conformational states and ligand binding contribute to stability in an interdependent manner. Our results are compatible with a scenario where ligand entry occurs in an open, partially unfolded state followed by subsequent structural compaction, completing a transition along Type I rearragement along a pathway including a different, Type II-like opening.

**Fig 5:**
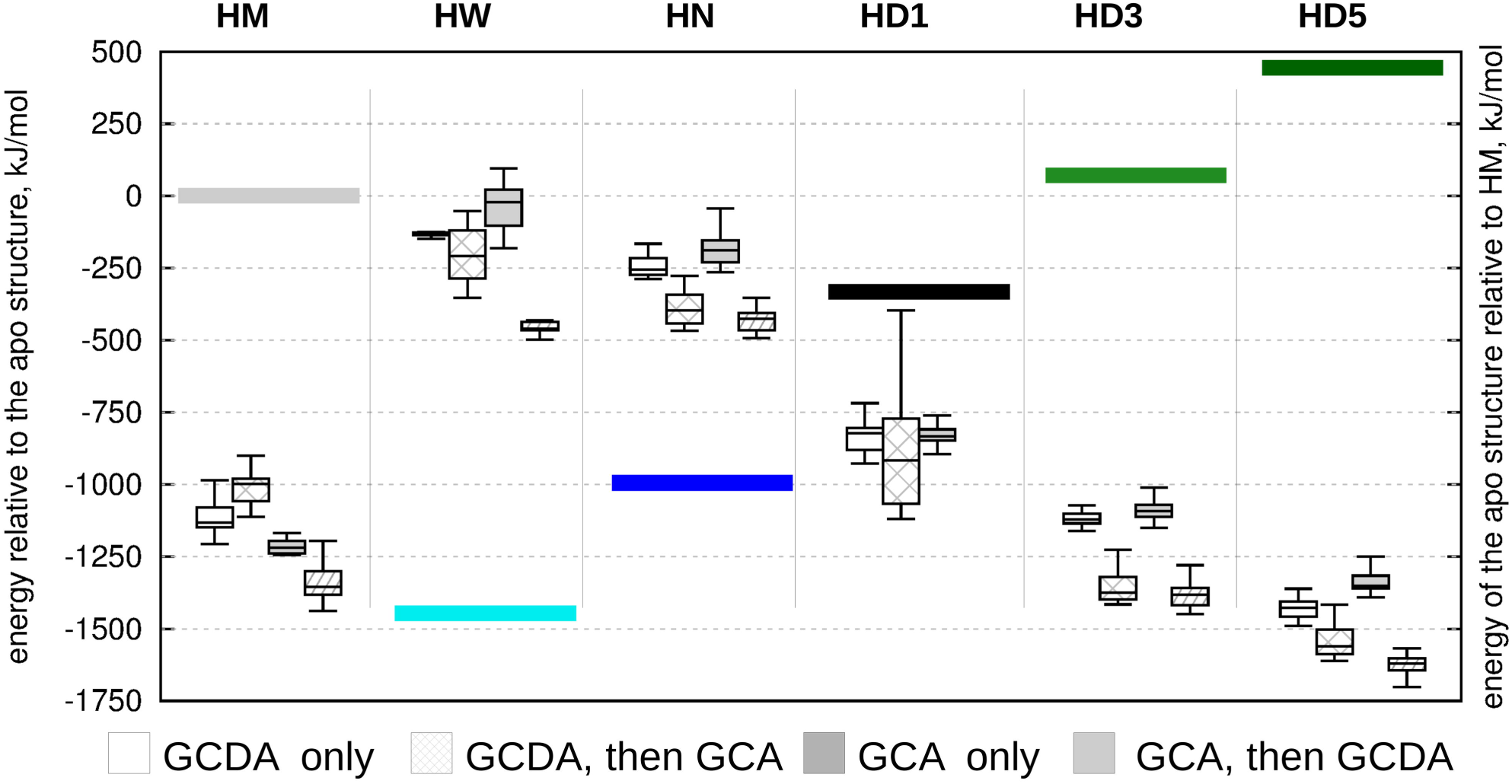
Relative energies of docked structures relative to the ligand-free conformations. Differences between the starting conformations of the molecular dynamics- derived structures relative to 2MM3 are depicted with colored bars. White boxes: only GCDA docked, stripped white boxes: GCDA docked first, GCA docked second, gray boxes: only GCDA docked, stripped gray boxes: GCA docked first, GCDA docked second. HM denotes a representative conformer from the holo MUMO esnemble, HN (holo narrow) and HW (holo wide) are selected extreme structures from the holo MUMO esnembles corresponding to Type II opening. In addition, three from the high correlation hidden conformers are selected (HD1, HD3 and HD5).

Comparison of ligand positions in the different structures (Table 3) and the total energies of the complexes (Fig 5) leads to the conclusion that open gastrotropin structures can bind ligands with a high structural versatility while maintaining high affinity. This suggests that ligands might undergo dynamic reposition even in the binary and ternary complexes.

**Table 3:**
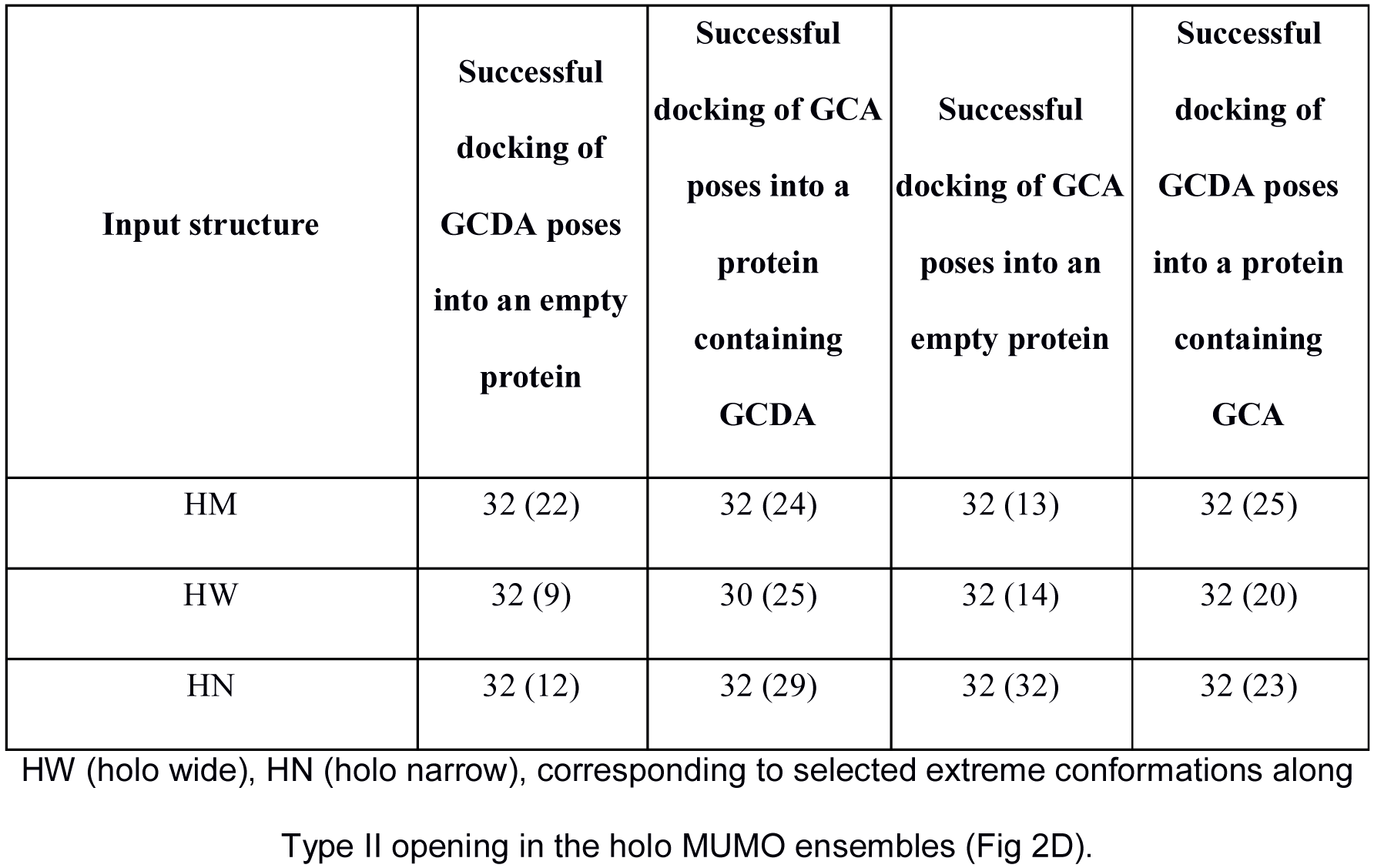
A summary of the docking simulations conducted on three specific input structures showing the number of successful calculations as well as the number of cases where the ligand binds in an orientation similar to that observed in the 2MM3 structure.

## Conclusions

We have generated structural ensembles that are in agreement with available NMR parameters reporting on the structure and fast time-scale dynamics of human gastrotropin. The two types of barrel opening identified are in agreement with previous observations of the iLBP family. We propose a refined model of ligand entry that is compatible with the portal hypothesis, namely, that the structural transition from the *apo* to the *holo* state, termed Type I opening, proceeds along an indirect route involving partial unfolding of the helical cap structure. In our model this unfolding is related to and facilitated by another mode of barrel opening, termed Type II, that is present in both the *apo* and *holo* states.

## Author contributions

ZG designed the study, ZH and ALS performed MD simulations, ALS performed docking calculations, all authors participated in data analysis and interpretation as well as in writing the manuscript.

## Supporting information

**S1 Fig. Experimental and back-calculated S2 order parameters parameters of different ensembles.**

(A) Apo MUMO ensembles by amino acids

(B) Apo MUMO ensembles as correlation plots

(C) Holo MUMO ensembles by amino acids

(D) Holo MUMO ensembles as correlation plots

(E) Unrestrained ensembles by amino acids

(F) Unrestrained ensembles as correlation plots

The PDB ensembles are depicted in all panels as references. All the plots were generated by the CoNSEnsX webserver.

**S2 Fig. Cα atom distance correlations.**

(A) Cα atom distance correlation matrix with PC1 (above diagonal) and PC2 (below diagonal) for each amino acid pair. Helical regions are labeled with beige rectangles, β strand regions with gray rectangles

(B) The highest (red, brown lines) and lowest (blue, gray lines) distances depicted on the structure correalted with PC1

(C) The lowest (blue, gray lines) distances depicted on the structure correalted with PC2.

**S3 Fig. Investigation of the docked frames.**

(A) Tanimoto distances of the simulation frames and the docked frames based on ligand contact data (see text)

(B) Comparison of the average of the number of ligand heavy atoms (vertical axis) being closer, than 4 Å to each amino acid (horizontal axis) of CHO (blue all frames, black docked frames).

(C). Comparison of the average of the number of ligand heavy atoms (vertical axis) being closer, than 4 Å to each amino acid (horizontal axis) of GCH: green simulated frames, red docked frames.

Correlations are listed on the top.

**S4 Fig. Histograms of PC2 (Type II motion) of the MUMO ensembles along PC1 and PC2.**

**S1 Table: The number of frames in each simulation, where the respective hydrogen bonds are present.**

**S1 Movie. The movement along the first PCA mode PC1.**

**S2 Movie. The movement along the second PCA mode PC2.**

**S1 File. Topology file for the apo 283 MUMO 6 ns simulation.**

**S2 File. Topology file for the apo 291 MUMO 6 ns simulation.**

**S3 File. Topology file for the apo 298 MUMO 6 ns simulation.**

**S4 File. Topology file for the apo 313 MUMO 6 ns simulation.**

**S5 File. Topology file for the holo 283 MUMO 6 ns simulation.**

**S6 File. Topology file for the holo 291 MUMO 6 ns simulation.**

**S7 File. Topology file for the holo 298 MUMO 6 ns simulation.**

**S8 File. Topology file for the holo 313 MUMO 6 ns simulation.**

**S1 Appendix. Main features of protein-ligand interactions are similar in the simulated and docked structures.**

## References

[1] Smathers RL, Petersen DR. The human fatty acid-binding protein family: evolutionary divergences and functions. Hum Genomics. 2011 Mar;5(3):170–191. doi:10.1186/1479-7364-5-3-170

[2] Horváth G, Király P, Tárkányi G, Toke O. Internal motions and exchange processes in human ileal bile acid binding protein as studied by backbone (15)N nuclear magnetic resonance spectroscopy. Biochemistry. 2012 Mar 6;51(9):1848–1861. doi:10.1021/bi201588q

[3] Lin MC, Kramer W, Wilson FA. Identification of cytosolic and microsomal bile acid-binding proteins in rat ileal enterocytes. J Biol Chem. 1990 Sep 5;265(25):14986–14995. PMID: 2394709

[4] Alrefai WA, Gill RK. Bile acid transporters: structure, function, regulation and pathophysiological implications. Pharm Res. 2007 Oct;24(10):1803–1823. doi:10.1007/s11095-007-9289-1

[5] Veerkamp JH, Maatman RG. Cytoplasmic fatty acid-binding proteins: their structure and genes. Prog Lipid Res. 1995;34(1):17–52. doi:10.1016/0163-7827(94)00005-7

[6] Sessler RJ, Noy N. A ligand-activated nuclear localization signal in cellular retinoic acid binding protein-II. Mol Cell. 2005 Apr 29;18(3):343–353. doi:10.1016/j.molcel.2005.03.026

[7] Ayers SD, Nedrow KL, Gillilan RE, Noy N. Continuous nucleocytoplasmic shuttling underlies transcriptional activation of PPARgamma by FABP4. Biochemistry. 2007 Jun 12;46(23):6744–6752. doi:10.1021/bi700047a

[8] Nakahara M, Furuya N, Takagaki K, Sugaya T, Hirota K, Fukamizu A, et al. Ileal bile acid- binding protein, functionally associated with the farnesoid X receptor or the ileal bile acid transporter, regulates bile acid activity in the small intestine. J Biol Chem. 2005 Dec 23;280(51):42283–42289. doi:10.1074/jbc.M507454200

[9] Tochtrop GP, Richter K, Tang C, Toner JJ, Covey DF, Cistola DP. Energetics by NMR: site-specific binding in a positively cooperative system. Proc Natl Acad Sc U S A. 2002 Feb 19;99(4):1847–1852. doi:10.1073/pnas.012379199

[10] Eliseo T, Ragona L, Catalano M, Assfalg M, Paci M, Zetta L, et al. Structural and dynamic determinants of ligand binding in the ternary complex of chicken liver bile acid binding protein with two bile salts revealed by NMR. Biochemistry. 2007 Nov 6;46(44):12557–12567. doi:10.1021/bi7013085

[11] Turpin ER, Fang HJ, Thomas NR, Hirst JD. Cooperativity and site selectivity in the ileal lipid binding protein. Biochemistry. 2013 Jul 9;52(27):4723–4733. doi:10.1021/bi400192w

[12] Kouvatsos N, Thurston V, Ball K, Oldham NJ, Thomas NR, Searle MS. Bile acid interactions with rabbit ileal lipid binding protein and an engineered helixless variant reveal novel ligand binding properties of a versatile beta-clam shell protein scaffold. J Mol Biol.2007 Aug 31;371(5):1365–1377. doi:10.1016/j.jmb.2007.06.024

[13] Kurz M, Brachvogel V, Matter H, Stengelin S, Thüring H, Kramer W. Insights into the bile acid transportation system: the human ileal lipid-binding protein-cholyltaurine complex and its comparison with homologous structures. Proteins. 2003 Feb 1;50(2):312–328. doi:10.1002/prot.10289

[14] Horváth G, Bencsura Á, Simon Á, Tochtrop GP, DeKoster GT, Covey DF, et al. Structural determinants of ligand binding in the ternary complex of human ileal bile acid binding protein with glycocholate and glycochenodeoxycholate obtained from solution NMR. FEBS J. 2016 Feb;283(3):541–555. doi:10.1111/febs.13610

[15] Banaszak L, Winter N, Xu Z, Bernlohr DA, Cowan S, Jones TA. Lipid-binding proteins: a family of fatty acid and retinoid transport proteins. Adv Protein Chem. 1994;45:89–151. doi:10.1016/S0065-3233(08)60639-7

[16] Tochtrop GP, Bruns JL, Tang C, Covey DF, Cistola DP. Steroid ring hydroxylation patterns govern cooperativity in human bile acid binding protein. Biochemistry. 2003 Oct 14;42(40):11561–11567. doi:10.1021/bi0346502

[17] Toke O, Monsey JD, DeKoster GT, Tochtrop GP, Tang C, Cistola DP. Determinants of cooperativity and site selectivity in human ileal bile acid binding protein. Biochemistry. 2006 Jan 24;45(3):727–737. doi:10.1021/bi051781p

[18] Tochtrop GP, DeKoster GT, Covey DF, Cistola DP. A single hydroxyl group governs ligand site selectivity in human ileal bile acid binding protein. J Am Chem Soc. 2004 Sep 8;126(35):11024–11029. doi:10.1021/ja047589c

[19] Horváth G, Egyed O, Toke O. Temperature dependence of backbone dynamics in human ileal bile acid-binding protein: implications for the mechanism of ligand binding. Biochemistry. 2014 Aug 12;53(31):5186–5198. doi:10.1021/bi500553f

[20] Hodsdon ME, Cistola DP. Discrete backbone disorder in the nuclear magnetic resonance structure of apo intestinal fatty acid-binding protein: implications for the mechanism of ligand entry. Biochemistry. 1997 Feb 11;36(6):1450–1460, doi:10.1021/bi961890r

[21] Jenkins AE, Hockenberry JA, Nguyen T, Bernlohr DA. Testing of the portal hypothesis: analysis of a V32G, F57G, K58G mutant of the fatty acid binding protein of the murine adipocyte. Biochemistry. 2002 Feb 12;41(6):2022–2027. doi:10.1021/bi015769i

[22] Ragona L, Pagano K, Tomaselli S, Favretto F, Ceccon A, Zanzoni S, et al. The role of dynamics in modulating ligand exchange in intracellular lipid binding proteins. Biochim Biophys Acta. 2014 Jul;1844(7):1268–1278. doi:10.1016/j.bbapap.2014.04.011.

[23] Cogliati C, Ragona L, D’Onofrio M, Günther U, Whittaker S, Ludwig C, et al. Site- specific investigation of the steady-state kinetics and dynamics of the multistep binding of bile acid molecules to a lipid carrier protein. Chemistry. 2010 Oct 4;16(37):11300–11310. doi:10.1002/chem.201000498

[24] Horváth G, Biczók L, Majer Z, Kovács M, Micsonai A, Kardos J, et al. Structural insight into a partially unfolded state preceding aggregation in an intracellular lipid-binding protein. FEBS J. 2017 Nov;284(21):3637–3661. doi:10.1111/febs.14264

[25] Ángyán AF, Gáspári Z. Ensemble-based interpretations of NMR structural data to describe protein internal dynamics. Molecules. 2013 Aug 30;18(9):10548–10567. doi:10.3390/molecules180910548

[26] Van Der Spoel D, Lindahl E, Hess B, Groenhof G, Mark AE, Berendsen HJ. GROMACS: fast, flexible and free. J Comput Chem. 2005 Dec;26(16):1701–1718. doi:10.1002/jcc.20291

[27] Pronk S, Páll S, Schulz R, Larsson P, Bjelkmar P, Apostolov R, et al. GROMACS 4.5: a high-throughput and highly parallel open source molecular simulation toolkit. Bioinformatics. 2013 Apr 1;29(7):845–854. doi:10.1093/bioinformatics/btt055

[28] Fizil Á, Gáspári Z, Barna T, Marx F, Batta G. “Invisible” conformers of an antifungal disulfide protein revealed by constrained cold and heat unfolding, CEST-NMR experiments and molecular dynamics calculations. Chemistry. 2015 Mar 23;21(13):5136–5144. doi:10.1002/chem.201404879

[29] Richter B, Gsponer J, Várnai P, Salvatella X, Vendruscolo M. The MUMO (minimal under-restraining minimal over-restraining) method for the determination of native state ensembles of proteins. J Biomol NMR. 2007 Feb;37(2):117–135. doi:10.1007/s10858-006-9117-7

[30] Kony D, Damm W, Stoll S, Van Gunsteren WF. An improved OPLS-AA force field for carbohydrates. J Comput Chem. 2002 Nov 30;23(15):1416–1429. doi:10.1002/jcc.10139

[31] Jorgensen WL, Chandrasekhar J, Madura JD, Impey RW, Klein ML. Comparison of simple potential functions for simulating liquid water. J Chem Phys. 1983 Apr;79(2):926–935. doi:10.1063/1.445869

[32] Ángyán AF, Szappanos B, Perczel A, Gáspári Z. CoNSEnsX: an ensemble view of protein structures and NMR-derived experimental data. BMC Struct Biol. 2010 Oct 29;10:39. doi:10.1186/1472-6807-10-39

[33] Hess B, Bekker H, Berendsen HJC, Fraaije JGEM. LINCS: A Linear Constraint Solver for Molecular Simulations. J Comput Chem. 07 Dec 1998;18(12):1463–1472. doi:10.1002/(SICI)1096-987X(199709)18:12<1463::AID-JCC4>3.0.CO;2-H

[34] Gabriel E, Fagg GE, Bosilca G, Angskun T, Dongarra JJ, Squyres JM, et al. Open MPI: Goals, Concept and Design of a Next Generation MPI Implementation. In: Kranzlmüller D, Kacsuk P, Dongarra J, editors. Recent Advances in Parallel Virtual Machine and Message Passing Interface. EuroPVM/MPI Lecture Notes in Computer Science, vol 3241. pp 97–104; 2004; Springer, Berlin, Heidelberg, Germany. doi:10.1007/978-3-540-30218-6_19

[35] Friesner RA, Murphy RB, Repasky MP, Frye LL, Greenwood JR, Halgren TA, et al. Extra precision glide: docking and scoring incorporating a model of hydrophobic enclosure for protein-ligand complexes. J Med Chem. 2006 Oct 19;49(21):6177–6196. doi:10.1021/jm051256o

[36] Dudola D, Kovács B, Gáspári Z. CoNSEnsX+ Webserver for the Analysis of Protein Structural Ensembles Reflecting Experimentally Determined Internal Dynamics. J Chem Inf Model. 2017 Aug 28;57(8):1728–1734. doi:10.1021/acs.jcim.7b00066

[37] Bakan A, Meireles LM, Bahar I. ProDy: Protein Dynamics Inferred from Theory and Experiments. Bioinformatics. 2011 Jun 1;27(11):1575–1577. doi:10.1093/bioinformatics/btr168

[38] Humphrey W, Dalke A, Schulten K. VMD: visual molecular dynamics. J Mol Graph. 1996 Feb;14(1):33–38. doi:10.1016/0263-7855(96)00018-5

[39] Han B, Liu Y, Ginzinger SW, Wishart DS. SHIFTX2: significantly improved protein chemical shift prediction. J Biomol NMR. 2011 May;50(1):43–57. doi:10.1007/s10858-011-9478-4

[40] Xu D, Tsai CJ, Nussinov R. Hydrogen bonds and salt bridges across protein-protein interfaces. Protein Eng Des Sel. 1997 Sep;10(9):999–1012. doi:10.1093/protein/10.9.999

[41] Baker EN, Hubbard RE. Hydrogen bonding in globular proteins. Prog Biophys Mol Biol. 1984;44(2):97–179. doi:10.1016/0079-6107(84)90007-5

[42] Cai J, Lücke C, Chen Z, Qiao Y, Klimtchuk E, Hamilton JA. Solution structure and backbone dynamics of human liver fatty acid binding protein: fatty acid binding revisited. Biophys J. 2012 Jun 6;102(11):2585–2594. doi:10.1016/j.bpj.2012.04.039

[43] Lücke C, Zhang F, Hamilton JA, Sacchettini JC, Rüterjans H. Solution structure of ileal lipid binding protein in complex with glycocholate. Eur J Biochem. 2000 May;267(10):2929– 2938. doi:10.1046/j.1432-1327.2000.01307.x

[44] Czajlik A, Kovács B, Permi P, Gáspári Z. Fine-tuning the extent and dynamics of binding cleft opening as a potential general regulatory mechanism in parvulin-type peptidyl prolyl isomerases. Sci Rep. 2017 Mar 16;7:44504. doi:10.1038/srep44504

[45] Pettersen EF, Goddard TD, Huang CC, Couch GS, Greenblatt DM, Meng EC, et al. UCSF Chimera-a visualization system for exploratory research and analysis. J Comput Chem. 2004 Oct;25(13):1605–1612. doi:10.1002/jcc.20084

[46] Andersen CA, Palmer AG, Brunak S, Rost B. Continuum secondary structure captures protein flexibility. Structure. 2002 Feb;10(2):175–184. doi:10.1016/S0969-2126(02)00700-1

